# Divergent patterns of meiotic double strand breaks and synapsis initiation dynamics suggest an evolutionary shift in the meiosis program between American and Australian marsupials

**DOI:** 10.1101/2023.02.13.527877

**Authors:** F. Javier Valero-Regalón, Mireia Solé, Pablo López-Jiménez, María Valerio-de Arana, Marta Martín-Ruiz, Roberto de la Fuente, Laia Marín-Gual, Marilyn B. Renfree, Geoff Shaw, Soledad Berríos, Raúl Fernández-Donoso, Paul D. Waters, Aurora Ruiz-Herrera, Rocío Gómez, Jesús Page

## Abstract

In eutherian mammals, hundreds of programmed DNA double-strand breaks (DSBs) are generated at the onset of meiosis. The DNA damage response is then triggered. Although the dynamics of this response is well studied in eutherian mammals, recent findings have revealed different patterns of DNA damage signaling and repair in marsupial mammals. To better characterize these differences, here we analyzed synapsis and the chromosomal distribution of meiotic DSBs markers in three different marsupial species (*Thylamys elegans, Dromiciops gliorides*, and *Macropus eugenii*) that represent South American and Australian Orders. Our results revealed inter-specific differences in the chromosomal distribution of DNA damage and repair proteins, which were associated with differing synapsis patterns. In the American species *T. elegans* and *D. gliroides*, synapsis progressed exclusively from the chromosomal ends towards interstitial regions. This was accompanied by sparse H2AX phosphorylation, mainly accumulating at chromosomal ends, which appeared conspicuously polarized in a *bouquet* configuration at early stages of prophase I. Accordingly, RAD51 and RPA were mainly localized at chromosomal ends throughout prophaseI in both American marsupials, likely resulting in reduced recombination rates at interstitial positions. In sharp contrast, synapsis initiated at both interstitial and distal chromosomal regions in the Australian representative *M. eugenii*, γH2AX had a broad nuclear distribution, and RAD51 and RPA foci displayed an even chromosomal distribution. Given the basal evolutionary position of *T. elegans*, it is likely that the meiotic features reported in this species represent an ancestral pattern in marsupials and that a shift in the meiotic program occurred after the split of *D. gliroides* and the Australian marsupial clade. Our results open intriguing questions about the regulation and homeostasis of meiotic DSBs in marsupials. The low recombination rates observed at the interstitial chromosomal regions in American marsupials can result in the formation of large linkage groups, thus having an impact in the evolution of their genomes.

## INTRODUCTION

Meiosis is a complex and highly regulated process, by which homologous chromosomes synapse, recombine and segregate. Synapsis refers to the tight association of homologs during meiotic prophase I by a structure called the synaptonemal complex (SC). The SC is formed by two axial/lateral elements (AE/LEs), one per homologue, held together by transverse filaments (TFs), which emanate from each of the LEs and overlap in a central region to form the central element (CE) (von Wettstein et al., 1984; Page and Hawley, 2004). Recognition of homologues in mammals (and many other organisms) is mediated by the formation of hundreds of programmed DNA double-strand breaks (DSBs) by the SPO11 protein at the beginning of prophase I (leptotene stage) (Keeney et al., 2014). The formation of DSBs triggers a DNA damage response that follows the homologous repair pathway, leading to the molecular interaction of chromosomes. The broken DNA molecule uses the intact DNA sequence of the homologue as a template for DNA repair during zygotene. These molecular interactions, in turn, stimulate and facilitate the synapsis of homologous chromosomes. In mammals (but not in other organisms), most DSBs produced during meiosis are repaired through a process that leads to gene conversion (non-reciprocal recombination events), whereas some of them result in reciprocal exchange events that lead to the formation of crossovers (COs) at the end of pachytene (at least one CO per bivalent) (Cole et al., 2012). These COs are visualized cytologically as chiasmata, which hold recombined homologous chromosomes together until they segregate during anaphase of the first meiotic division (Roeder, 1997).

Besides a role in ensuring faithful chromosome segregation, it is commonly accepted that recombination increases genetic variability in natural populations through the generation of new haplotypes, which are later subjected to evolutionary drift and selection (Barton and Charlesworth, 1998; Otto and Lenormand, 2002). In contrast, supression of recombination at specific chromosomal regions leads to the genetic isolation of these chromosome segments and the formation of large linkage groups. If allele combinations cannot be reshuffled by recombination, beneficial alleles are likely to be lost by either background selection or random drift (Graves, 1995; Charlesworth et al., 2005; Bachtrog, 2013). Finally, both gene conversion and CO formation can alter the GC content of genomic regions where these events accumulate (called hotspots) by a process known as GC-biased gene conversion (gBGC) (Duret and Galtier, 2009). Therefore, the frequency and distribution of meiotic recombination have a significant impact on genome evolution (Lenormand et al., 2016; Bergero et al., 2021).

In mammals, meiotic studies have been traditionally focused in model species, mainly the house mouse and humans. However, comparative studies are important to understand if the features described in these models are present in other species. For instance, the organization and composition of the SC seem to be particularly well conserved (Fraune et al., 2012). Other features, like the frequency of recombination, have also received great attention, though they are more variable between species (Dumont and Payseur, 2008; Segura et al., 2013). Additional aspects, like the regulation of chromosome segregation, remain unexplored in most mammals, especially in non-eutherians. This is the case in marsupials, the sister group of eutherian mammals, which diverged from each other around 165 million years ago. There are currently about 270 marsupial species, distributed in America and Australia. They are grouped into two main clades: Ameridelphia, which comprises the Orders Didelphimorphia and Paucituberculata; and Australidelphia, which includes the Australian Orders Dasyuromorphia, Peramelemorphia, Notoryctemorphia and Diprodontia (Figure 1A) (Duchêne et al., 2017). Intriguingly, Australidelphia also includes an American sister clade, the Order Microbiotheria, only represented by two species of monito del monte (*Dromiciops gliroides* and *D. vozinovici*) (D’Elía et al., 2016; Feng et al., 2022; Fontúrbel et al., 2022).

**FIGURE 1.**
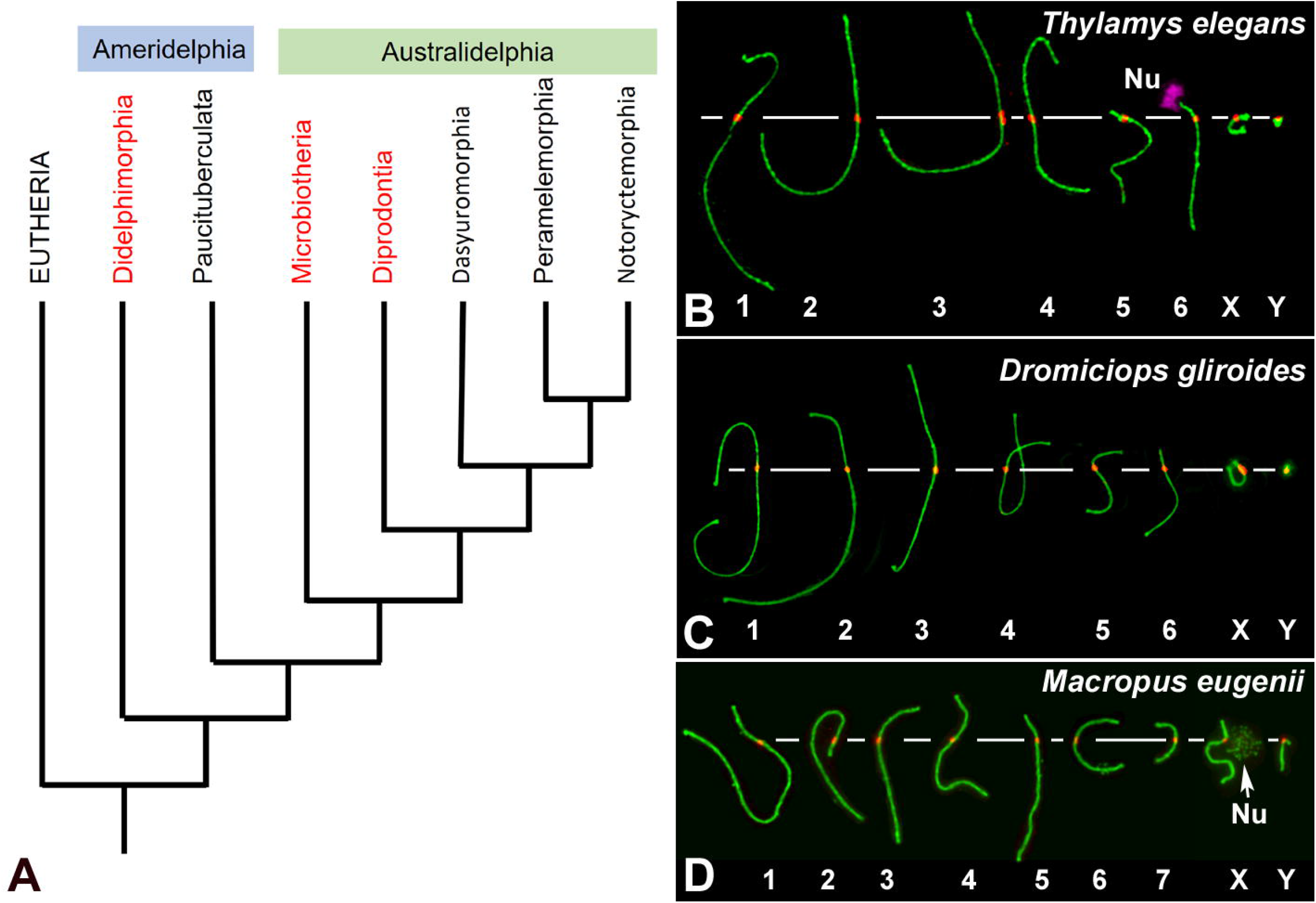
**A**. Phylogenetic relationships of extant marsupial orders. The arrangement presented is based on the phylogeny published by Duchêne and coworkers (Duchêne et al., 2017). Although the topology of the tree is still controversial, Microbiotheria is grouped to Australidelphia in all the trees consulted. The Orders included in this study are highlighted in red. **B-D**. Meiotic karyotypes of the three studied species: SYCP3 in green and centromeres in red. Bivalents are ordered by size, according to previous reports. In *T. elegans* (**B**) the position of the NOR (Nu) on the short arm of bivalent 6 was detected using ani-fibrillarin antibody (pink). The position of the NOR on the X chromosome of *M. eugenii* was detected by an accumulation of SYCP3.

Marsupials are characterized by their unique reproductive strategy, in which pregnancy is uniformly short and the altricial young are born at an early developmental stage. Development is usually completed within an abdominal pouch, with the pouch young dependent on a highly specialized milk (Tyndale-Biscoe and Renfree, 1987). Marsupials also present a number of genetic and chromosomal differences compared to eutherians (Graves and Renfree, 2013). Two of the most relevant are: 1) Their reduced number of chromosomes (Hayman, 1990; Deakin, 2018). Although chromosome numbers range from 2n=10 to 2n=34, they present a bimodal distribution and most species have either 2n=14 or 2n=24 (Deakin and Potter, 2019; Deakin and O’Neill, 2020). Since the genome size is comparable to that of eutherians, marsupial chromosomes are usually much larger. 2) The Y chromosome is generally tiny and does not share a pseudoautosomal region (PAR) with the X due to extreme degeneration of the Y over evolutionary time (Graves et al., 1998). In fact, the Y chromosome can be lost in some somatic tissues, as reported in males of the family Peramelidae (Watson et al., 1998).

The special features of marsupial chromosomes also have an impact on their behavior during meiosis. The most noticeable feature is the behavior of sex chromosomes. The absence of a PAR on the XY pair precludes their reciprocal synapsis and recombination in male meiosis, thus challenging the usual way by which homologous chromosomes ensure their segregation during first meiotic division. Instead, sex chromosomes in marsupials present an alternative mode of association, which relies on the formation of a specific structure called the dense plate (DP) that maintains the sex chromosome association from prophase I (Solari and Bianchi, 1975; Sharp, 1982; Seluja et al., 1987; Page et al., 2003) until they segregate at anaphase-I (Page et al., 2006). Although small differences between species have been found regarding the timing of DP formation (Marín-Gual et al., 2022b), the development and dynamics of the DP are well conserved (Sharp, 1982; Page et al., 2005; Fernández-Donoso et al., 2010), indicating that the DP represents a feature that originated before the radiation of marsupials (Page et al., 2005). The emergence of this alternative mechanism of segregation allowed for the proper transmission of sex chromosomes after their complete differentiation. Interestingly, analogous mechanisms of sex chromosome segregation have independently appeared in eutherian species with completely differentiated sex chromosomes (de la Fuente et al., 2007; de la Fuente et al., 2012; Gil-Fernández et al., 2020; Gil-Fernández et al., 2021).

The striking behavior of sex chromosomes may have obscured other meiotic differences in marsupials. This includes recombination rates, i.e., the number of COs per cell, which are lower in marsupials compared to eutherians (Zenger et al., 2002; Samollow et al., 2004; Samollow et al., 2007; Dumont and Payseur, 2008; Wang et al., 2011). Moreover, marsupial males seem to be more recombinogenic than females, as opposed to the higher recombination rates in most female eutherian mammals (Bennett et al., 1986; Samollow et al., 2004; Samollow et al., 2007). We have recently proposed that the low recombination rates observed in marsupial males might result from the induction of fewer DSBs during prophase I, potentially leading to the formation of fewer COs (Marín-Gual et al., 2022b). Our previous study revealed that in three species of phylogenetically distant marsupials, the overall number of DSBs was significantly lower than in eutherian mammals (i.e., mice and humans), concomitant with low γH2AX levels on autosomes. Moreover, we detected inter-specific differences in the pattern of DSBs distribution along chromosomes (Marín-Gual et al., 2022b). Among eutherians, DSBs tend to accumulate slightly towards the distal regions of chromosomes in mice (Pratto et al., 2014), although COs show a more even distribution (Froenicke et al., 2002). The polarization of DSBs and COs towards the chromosomal ends is more evident in humans (Oliver-Bonet et al., 2007).

Here we test whether the distribution of DSBs in marsupials is even along chromosomes. To achieve this, we analyzed the localization of proteins related to DNA damage response and repair (γH2AX, RAD51 and RPA), along with SC components (SYCP1 and SYCP3), during meiosis in species that capture the deepest divergences within marsupials: the American species *Thylamys elegans* and *Dromiciops gliroides* and the Australian species *Macropus eugenii*. Our results uncover remarkable differences in the initiation and progression of synapsis between homologous chromosomes, as well as in the distribution pattern of DNA repair markers, with American species showing an extreme polarization towards chromosomal ends. This behavior may have important consequences for recombination rates and distribution, which in turn could impact genome evolution.

## MATERIALS AND METHODS

### Animals

Two *Thylamys elegans* (Didelphidae) and two *Dromiciops gliroides* (Microbiotheriidae) males were collected in central and Southern Chile, respectively, from natural populations under permission of Corporación Nacional Forestal (Conaf). Handling of animals was performed according to the ethical rules stablished by the University of Chile. Two *Macropus eugenii* (Macropodidae) males were collected from wild populations originating on Kangaroo Island (South Australia) that were later held in a breeding colony in Melbourne (Victoria, Australia). Sampling was conducted under ethics approval from the University of Melbourne Animal Experimentation Ethics Committees and followed the Australian National Health and Medical Research (2013) guidelines. The karyotypes of these species are as follows: *T. elegans* 2n=14; *D. gliroides* 2n=14; *M. eugenii* 2n=16. The meiotic karyotypes of the three species were arranged according to length and centromere position of each bivalent (Figure 1B-D), in agreement with previous reports (Page et al., 2003; Fernández-Donoso et al., 2010; Marín-Gual et al., 2022b).

### Spermatocyte spreads and squashes

Testicular samples were obtained and subsequently processed. For spreads, we used the protocol previously described by Peters and coworkers (Peters et al., 1997), with slight modifications for marsupial samples (Page et al., 2005). Briefly, a cell suspension was incubated in 10 mM sucrose solution in distilled water for 15 minutes. The suspension was spread onto a slide dipped in 1% formaldehyde in distilled water (pH 9.5), containing 100 mM sodium tetraborate and 0.15% Triton-X100. Cells were left to settle for 1.5 hours in a humid chamber and subsequently washed with 0.4% Photoflo (Kodak) in distilled water. Slides were air dried at room temperature and then rehydrated in phosphate saline buffered (PBS: NaCl 137 mM, KCl 2,7 mM, Na2HPO4 10,1 mM, KH2PO4 1,7 mM, pH 7.4) before immunostaining. For squashes, we used a previously described method (Page et al., 1998; Page et al., 2003). Seminiferous tubules were fixed in 2% formaldehyde in PBS for 10 minutes and then squashed on a slide. Coverslip was removed after freezing in liquid nitrogen and slides were rehydrated in PBS until use.

### Immunofluorescence

Slides were incubated overnight at 4°C with the following antibodies diluted in PBS: rabbit anti-SYCP3 (ab15093, Abcam, 1:200 dilution), rabbit anti-SYCP1 (ab15087, Abcam, 1:200 dilution), mouse anti-γH2AX (05-636, Upstate, 1:1000 dilution), rabbit anti-RAD51 (PC130, Calbiochem, 1:50 dilution), rabbit anti-RPA2 (ab10359, Abcam, 1:50 dilution), mouse anti-fibrillarin (ab4566, Abcam; 1:50 dilution), human anti-centromere (441-10BK-50, Antibodies Incorporated, 1:50 dilution). In addition, many antibodies were used against DMC1, MLH1, MLH3, and other proteins associated with COs (PRR19, CNTD1, CDK2) that yielded no positive labeling. After incubation, slides were washed three times in PBS and subsequently incubated for one hour at room temperature with secondary antibodies conjugated with Alexafluor 350, Alexafluor 488, Cy3 or Cy5 (Jackson ImmunoResearch Laboratories) all of them diluted 1:100 in PBS. After three washes in PBS slides were stained with 10 μg/ml DAPI, washed in PBS and mounted with Vectashield.

### Microscopy and image processing

Observations were made on an Olympus BX61 microscope equipped with appropriate fluorescence filters and an Olympus DP72 digital camera. The images were processed using the public domain software ImageJ (National Institutes of Health, USA; http://rsb.info.nih.gov/ij) and Adobe Photoshop 7.0 (Adobe). Spread images were taken as single-plane pictures, whereas squashed spermatocytes were photographed at 0.2 μm intervals and the resulting stack images processed in ImageJ.

### Quantitative analysis of RPA distribution

For the analysis of RPA foci chromosomal distribution, bivalents were identified at early pachytene spermatocytes according to their length and centromere position. In the case *T. elegans*, the location of fibrillarin signal associated to the short arm allowed the identification of bivalent 6. Each bivalent was measured using the *Free Hand* tool in ImageJ. The distance of centromeres and RPA foci from the tip of the short arm of the bivalents was assessed as follows: each focus was manually drawn as an intersection line with the outline of the SC, yielding the longitudinal position of the focus. Then, each bivalent was divided into 10 different segments, being segment 1 the distal portion of the shortest arm. Finally, the position of each RPA focus was ascribed to a specific segment (from 1 to 10). A minimum of 15 spermatocytes were recorded for each individual (two *T. elegans* and two *M. eugenii* males).

### Statistical analyses

Quantitative data were analyzed using Prism GraphPad 7.0. The distribution of RPA foci along chromosomes was compared to a random distribution by a χ^2^ goodness of fit test with 9 degrees of freedom. Statistical significance was considered for p<0.05. The relationship between RPA foci number and SC length was evaluated by Spearman correlation coefficient (r).

## RESULTS

### Chromosome synapsis dynamics

We first studied the synaptic behavior of chromosomes during meiosis in the selected species via the immunolocalization of the proteins SYCP3 and SYCP1, the main components of the axial/lateral elements (AE/LEs) and transverse filaments of the SC, respectively. The localization patterns of these proteins were used to classify spermatocytes into the different prophase I stages, following previous observations in marsupials (Page et al., 2003; Page et al., 2005; Marín-Gual et al., 2022b).

In *T. elegans*, during early prophase I SYCP3 was usually accompanied by the appearance of a SYCP1 signal (Figure 2A), making it difficult to discriminate between leptotene and zygotene. This suggests that the formation of the AEs was concurrent with the initiation of synapsis early in prophase I in this species. Thus, at early stages of prophase I the AEs were just partially formed, appearing with dotted signal along most of the chromosome, but forming short lines at the regions where two AEs associate (Figure 2A). These synapsed segments were mostly grouped in a small region (i.e. a *bouquet* configuration). In addition, SYCP3 revealed a thickening at the ends of the AEs. Therefore, we refer to this stage as the leptotene-zygotene transition. At a subsequent stage, early zygotene (Figure 2B), the AEs were almost completely formed. Synapsis began at chromosomal ends, which was evidenced by the presence of SYCP1 in the region where the AEs (now called LEs) of homologous chromosomes were associated. Moreover, the ends of chromosomes were still polarized in a *bouquet* conformation at this stage. SYCP1 was observed as continuous lines that regularly expanded from the ends towards the centers of the chromosomes, and there was no interstitial initiation of synapsis. This feature was still observable at late zygotene (Figure 2C). The only exception to synapsis beginning from telomeres was for the chromosome pair bearing the nucleolar organizing region (NOR) (see Figure 3C). The NOR is located near the telomere of the short arm (Figure 1) and it had delayed synapsis. Even though synapsis in the autosomes was completed by pachytene, the sex chromosomes remained unsynapsed at this stage (Figure 2D). In the other American species, *D. gliroides*, the pattern of AE formation and synapsis progression was almost identical, including the conspicuous *bouquet* configuration, the thickening of the distal regions of the LEs at early zygotene, the distal initiation of synapsis, and its subsequent progression to interstitial regions (Figure 2E-H).

**FIGURE 2.**
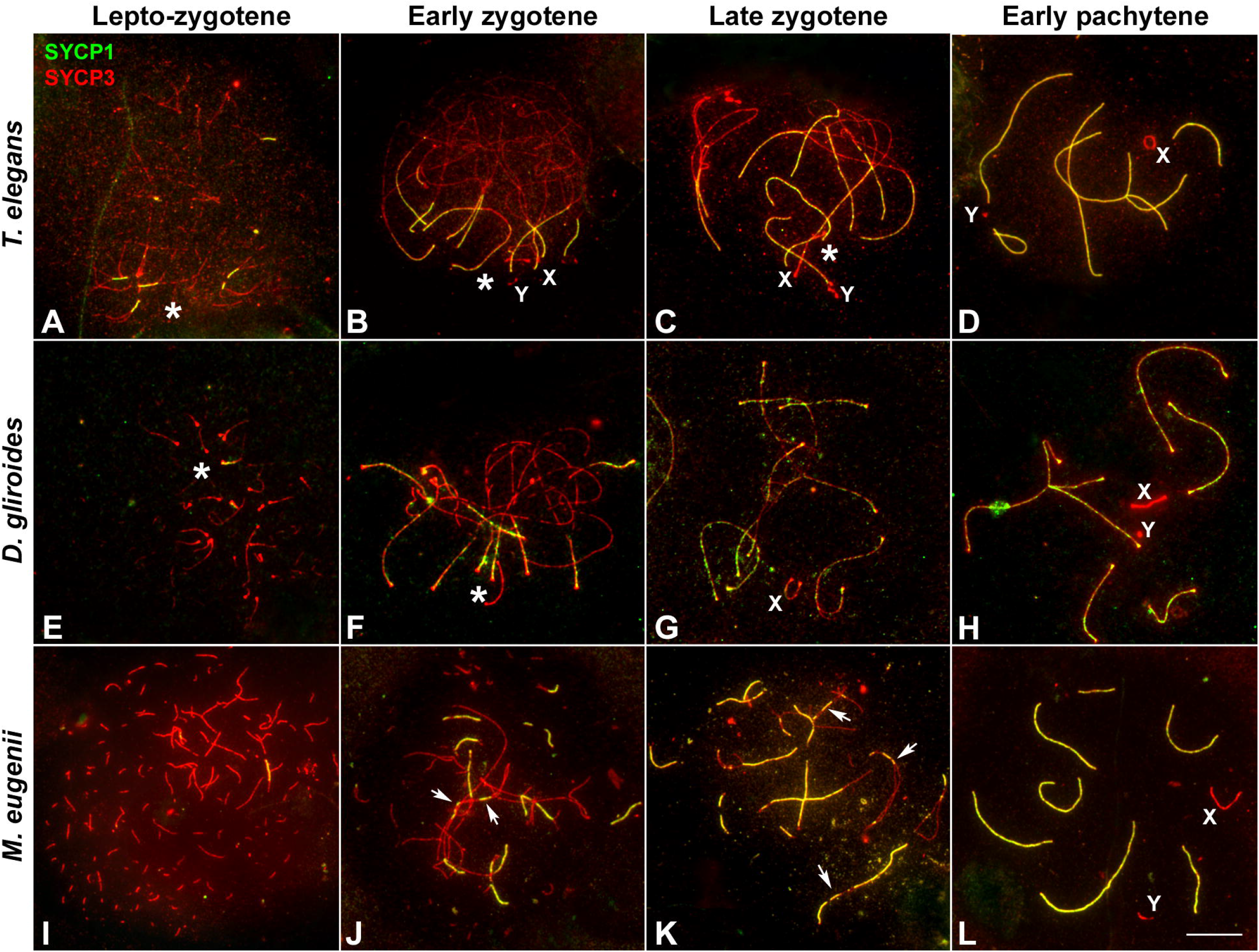
Synapsis progression during prophase I. Spread spermatocytes labeled with antibodies against SYCP3 (red) and SYCP1 (green). **A-D**. *T. elegans*. Short filaments of SYCP1 are seen between AEs at the leptotene-zygotene transition (**A**). These filaments appear mostly polarized to a specific nuclear region, the *bouquet* area (asterisk). Synapsis progresses during early (**B**) and late zygotene (**C**). Polarization of chromosomal ends is still observed (asterisks). Sex chromosomes (X, Y) lie in the *bouquet* region. Synapsis is complete at pachytene (**D**) except for the sex chromosomes. **E-H**. *D. gliroides*. Chromosomal ends are polarized to the *bouquet* area (asterisks) at leptotene-zygotene transition (**E**) and early zygotene (**F**). Synapsis progresses during late zygotene (**G**) and is complete at pachytene (**H**). Sex chromosomes remain separated. (**I-L**) *M. eugenii*. AEs appear as short fragments in the whole nucleus during the leptotene-zygotene transition (**I**). At early zygotene (**J**) synapsis is initiated both at the chromosomes ends and at interstitial regions (arrows). This is also observed at late zygotene (**K**). Synapsis is complete at pachytene (**L**), with sex chromosomes lying separately. Bar: 10 μm.

Remarkably, AE formation and SC assembly in *M. eugenii* contrasted the pattern in American species. At the leptotene-zygotene transition, the AEs appeared as short fragments or dots evenly distributed (Figure 2I). Some small fragments of SYCP1 signal were observed, but they did not adopt a polarized distribution. At early zygotene, the AEs appeared more continuous, and synapsis initiated both at the distal and interstitial regions of each chromosome (Figure 2J), a feature that was still detectable at late zygotene (Figure 2K). At pachytene, synapsis of the autosomes was complete, whereas sex chromosomes remained unsynapsed (Figure 2L).

### Distribution of DSBs during prophase I

In mammals, synapsis initiation is dependent on the occurrence of DNA DSBs at the beginning of meiosis (Baudat et al., 2000; Romanienko and Camerini-Otero, 2000). To assess if the differences detected in the progression of synapsis could be linked to a differential distribution of DSBs, we studied the localization of the phosphorylated form of histone H2AX (γH2AX), a widely used marker of DNA damage during meiosis (Mahadevaiah et al., 2001; Turner et al., 2004). Mirroring previous observation (Marín-Gual et al., 2022b) we found that in *T. elegans* only a few small foci of γH2AX became detectable at leptotene-zygotene on the chromatin around the AEs formation (Figure 3A). This location followed the pattern of chromosome synapsis described above, corresponding with chromosomal ends polarized in the *bouquet* configuration. At early zygotene, γH2AX labeling was mostly associated with the chromosomal regions where synapsis was initiated, whereas the rest of the nucleus remained devoid of γH2AX (Figure 3B). At this stage, the *bouquet* polarization was still observed. At late zygotene, an increase of γH2AX signal was observed, localized mainly over the regions of autosomes that had not completed synapsis, as well as over the chromatin around the AEs of the sex chromosomes (Figure 3C). At pachytene, once autosomes had completed full synapsis, γH2AX signal was only detectable over the sex chromosomes (Figure 3D). The distribution of γH2AX during meiosis in *D. gliroides* was very similar to that of *T. elegans*, i.e., γH2AX was only detected at the chromosomal ends at leptotene and early zygotene (Figure 3E-F). The phosphorylated histone accumulated at unsynapsed chromosomes in late zygotene (Figure 3G) and remained on sex chromosomes during pachytene (Figure 3H).

**FIGURE 3.**
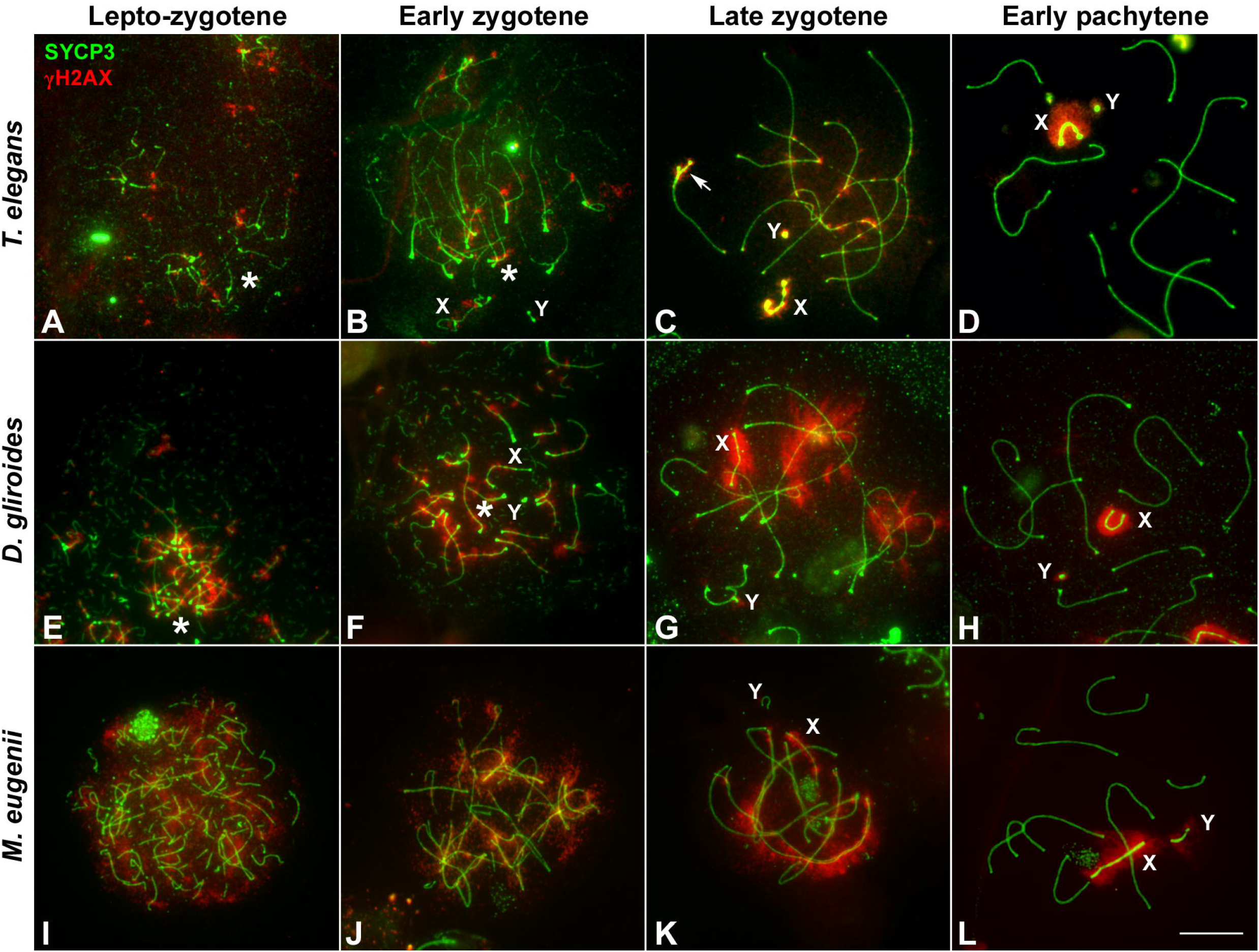
Localization of DNA damage-related proteins. Spread spermatocytes labeled with antibodies against SYCP3 (green) and γH2AX (red). **A-D**. *T. elegans*. γH2AX is observed as small foci at the leptotene-zygotene transition (**A**) and early zygotene (**B**), associated to the synapsing chromosomal ends polarized to the *bouquet* (asterisks). A more intense γH2AX labeling is observed at late zygotene (**C**) associated to the AEs of unsynapsed autosomal regions and sex chromosomes (X, Y). The proximal end of chromosome 6 also remains unsynapsed (arrow). At pachytene (**D**) γH2AX is only observed on the chromatin of sex chromosomes. **E-H**. *D. gliroides*. The pattern of γH2AX is almost identical to the one described for *T. elegans*, except for a more intense labeling of γH2AX in the unsynapsed regions of chromosomes at zygotene. (**I-L**) *M. eugenii*. At the leptotene to zygotene transition (**I**) γH2AX is spread in the whole nucleus. As zygotene proceeds (**J** and **K**) γH2AX labeling is reduced in the nucleus and concentrates around the AEs of unsynapsed autosomes and the X chromosome, but not on the Y chromosome. Both sex chromosomes exhibit signal of the antibody at pachytene (**L**). Bar: 10 μm.

Crucially, in *M. eugenii* the γH2AX signal was different. At the leptotene-zygotene transition (Figure 3I) the signal was distributed over all chromosomes, with no specific accumulations at chromosomal ends or any other region. This broad distribution was also observed at early zygotene (Figure 3J). At late zygotene, γH2AX tends to disappear from synapsed chromosomes but an intense signal was detected over the still unsynapsed autosomal regions and over the sex chromosomes (Figure 3K). At pachytene, γH2AX signal remained only over the sex chromosomes (Figure 3L).

The striking differences in the intensity and distribution of γH2AX between marsupial species lead us to test whether the faint signal observed in *T. elegans* could be due to an artifact of the spreading technique. Thus, we evaluated in this species the distribution of γH2AX in spermatocyte squashes, which maintained the three-dimensional organization of the nucleus and provided better preservation of the chromatin (Figure 4). This confirmed the patterns detected in spermatocyte spreads; that is, the almost complete absence of γH2AX at leptotene and early zygotene was a *bona fide* feature of *T. elegans* (Figure 4A-C). The accumulation of γH2AX at the unsynapsed regions started when synapsis had greatly progressed on the autosomes (Figure 4D-E). During pachytene, γH2AX was only present on the sex chromosomes, either before they paired (Figure 4F-G) or after they completed their pairing and the formation of the dense plate (DP) (Figure 4H).

**FIGURE 4.**
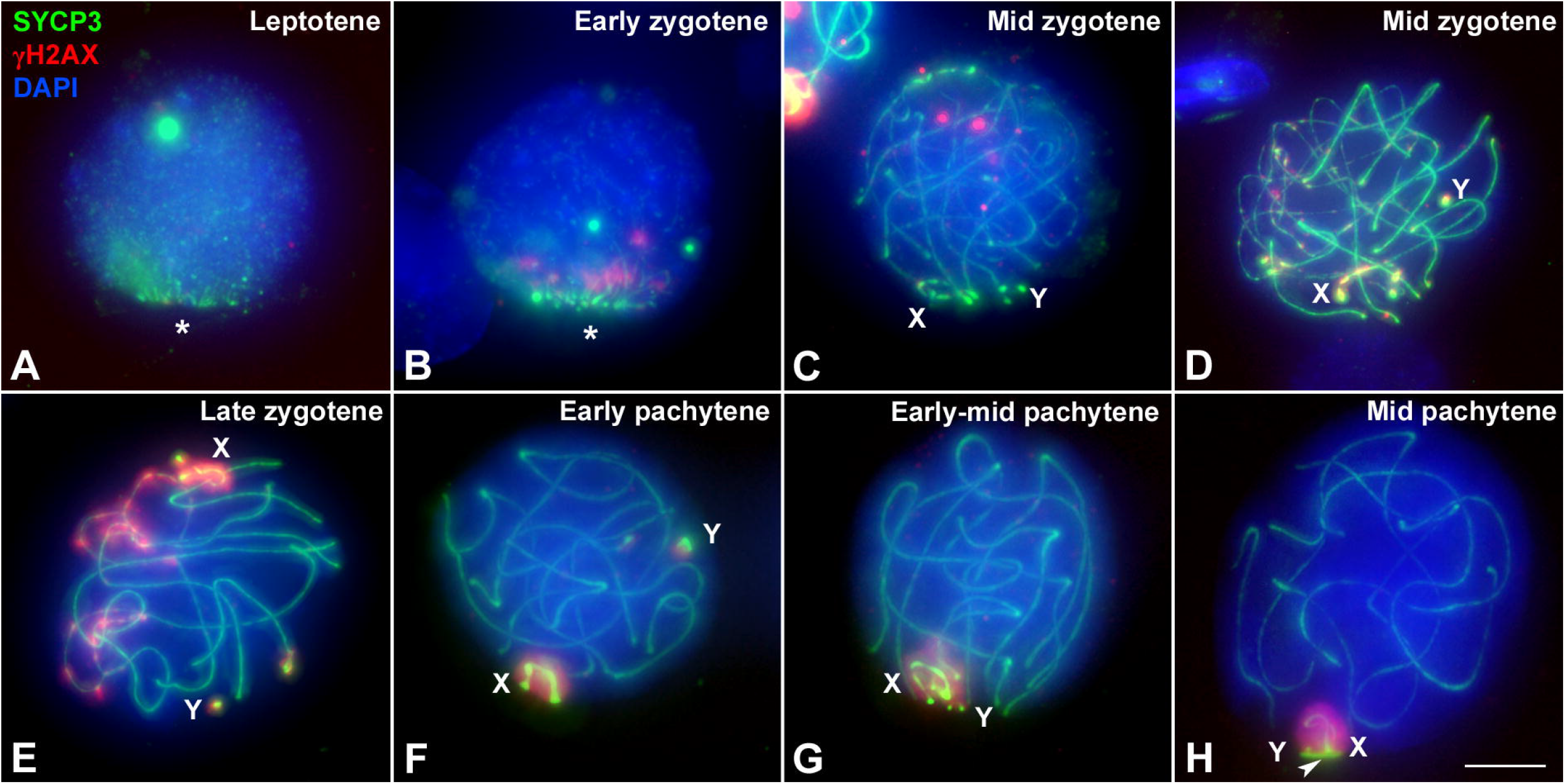
Localization of DNA damage in *T. elegans* spermatocytes preserving the 3-dimensional topology of chromosomes. Squashed spermatocytes labeled with antibodies against SYCP3 (green) and γH2AX (red) and DAPI (blue). **A**. Leptotene. No γH2AX labeling is observed. AEs appear polarized in a *bouquet* configuration (asterisk). **B**. Early zygotene. A few small γH2AX signals are observed at the region where synapsis is initiating. The *bouquet* polarization is still evident. **C**. Mid zygotene. Synapsis has progressed. A few γH2AX foci are scattered within the nucleus. **D**. Mid zygotene. γH2AX labeling starts to be observed at the unsynapsed regions of autosomes and sex chromosomes (X, Y). **E**. Late zygotene. γH2AX signal increases on unsynapsed chromosomes. Sex chromosomes are detected at opposite nuclear spaces and the γH2AX labeling extends from their AEs to the surrounding chromatin. **F**. Early pachytene. γH2AX is only detected on the chromatin of sex chromosomes, clearly separated in the nucleus. **G**. Early-mid pachytene. Sex chromosomes approach and associate to each other. **H**. Mid pachytene. Sex chromosomes pair and form the dense plate (arrowhead). γH2AX labels the whole sex body. Bar: 5 μm.

### Nuclear distribution of DNA repair proteins

The induction of DSBs triggers the activation of the homologous recombination repair pathway and the incorporation of proteins involved in this process, such as RAD51 and DMC1, the recombinases that mediate the invasion of an intact DNA template to repair the DSBs, and RPA, which protects single stranded DNA molecules generated during homologous recombination (Brown and Bishop, 2015). Here we report the chromosomal distribution of RAD51 and RPA in the species studied. Unfortunately, DMC1 did not yield a positive result.

We first analyzed the distribution of RAD51 in squashed spermatocytes. At the leptotene-zygotene transition, a few RAD51 foci were observed in *T. elegans*, mainly located at chromosomal ends and grouped in the *bouquet* configuration (Figure 5A). Some additional foci were observed scattered over the nucleus. At early zygotene (Figure 5B), foci remained localized mostly close to the chromosomal ends. At late zygotene (Figure 5C), the *bouquet* configuration was lost, and some RAD51 foci were localized interstitially along bivalents. The X chromosome accumulated numerous RAD51 foci at late zygotene and also at early pachytene (Figure 5D). A similar trend was observed for *D. gliroides* in spread spermatocytes (Figure 5E-H). Most foci were associated with the short SYCP3 filaments at leptotene (Figure 5E). Along with synapsis progression, some RAD51 foci appeared on interstitial regions of chromosomes (Figure 5F-G). This suggested progressive incorporation of RAD51 along chromosomes during zygotene. Some foci were still detectable at early pachytene (Figure 5H). Similar to *T. elegans*, the X chromosome presented abundant RAD51 foci (Figure 5H).

**FIGURE 5.**
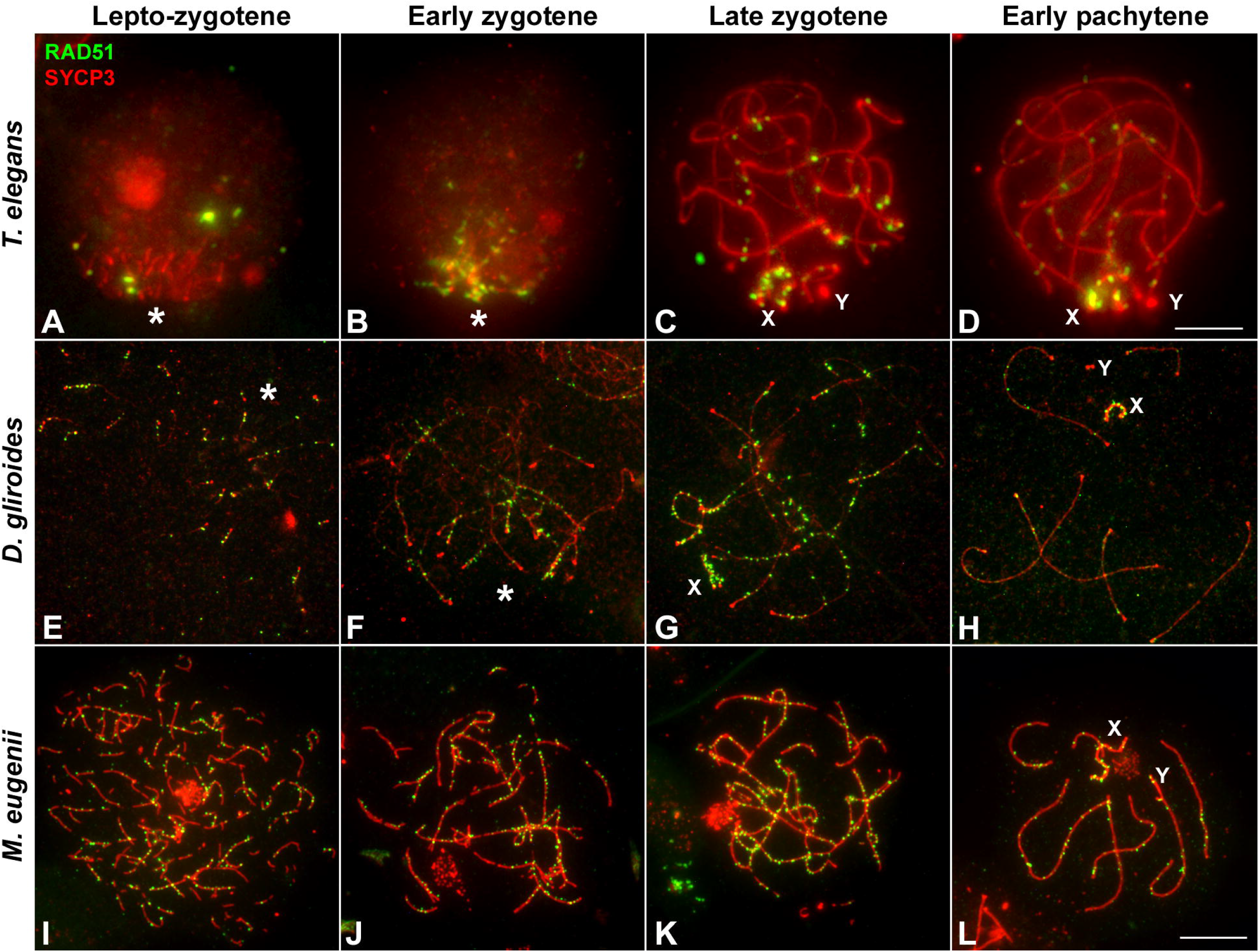
Localization of homologous recombination repair. Spermatocytes labeled with antibodies against SYCP3 (red) and RAD51 (green). **A-D**. *T. elegans*. Squashed spermatocytes. RAD51 foci are scarce and localized in the *bouquet* (asterisk) during the leptotene-zygotene transition (**A**) and also at early zygotene (**B**). At late zygotene (**C**) some RAD51 foci are seen at interstitial regions and coat abundantly the X and Y chromosomes (X, Y). At early pachytene (**D**) most RAD51 foci concentrate in the sex chromosomes. **E-H**. *D. gliroides*. RAD51 foci associate to the short stretches of SYCP3 at the leptotene-zygotene transition (**E**). At early zygotene (**F**), most foci are localized in the already synapsed distal regions. At late zygotene (**G**) discrete foci are also detected over the interstitial regions of autosomes and mostly over the X chromosome. At early pachytene (**H**), a few foci are still associated to autosomes. (**I-L**) *M. eugenii*. Spread spermatocytes. At the leptotene-zygotene transition (**I**) RAD51 foci appear in the whole nucleus. As zygotene proceeds (**J** and **K**) RAD51 is clearly observed all along the bivalents. At early pachytene (**L**), RAD51 foci number has decreased but they are observed all along autosomal bivalents and the sex chromosomes. Bars: 5 μm in **A-D** and 10 μm in **E-L**.

In sharp contrast, RAD51 foci appeared evenly distributed over the nucleus in *M. eugenii* spermatocyte spreads. At the leptotene-zygotene transition, foci were associated with the short fragments of the forming AEs (Figure 5I). Similarly, from early to late zygotene, RAD51 foci were distributed all along the synapsing bivalents (Figure 5J, K). Even at early pachytene, RAD51 foci did not concentrate at any particular chromosomal region, even though the number of such foci was prominently reduced.

As for RPA, we only obtained a reliable signal of the antibody in *T. elegans* and *M. eugenii*. The dynamics of RPA foci were similar to that of RAD51. In *T. elegans*, most foci were localized to the chromosomal ends at the leptotene-zygotene transition and early zygotene (Figure 6A, B). Then, foci also appeared at interstitial regions during late zygotene and early pachytene (Figure 5C and Figure 6D) but remained visibly concentrated at chromosomal ends. In contrast, RPA foci in *M. eugenii* were evenly distributed along chromosomes throughout prophase I (Figure 6E-H).

**FIGURE 6.**
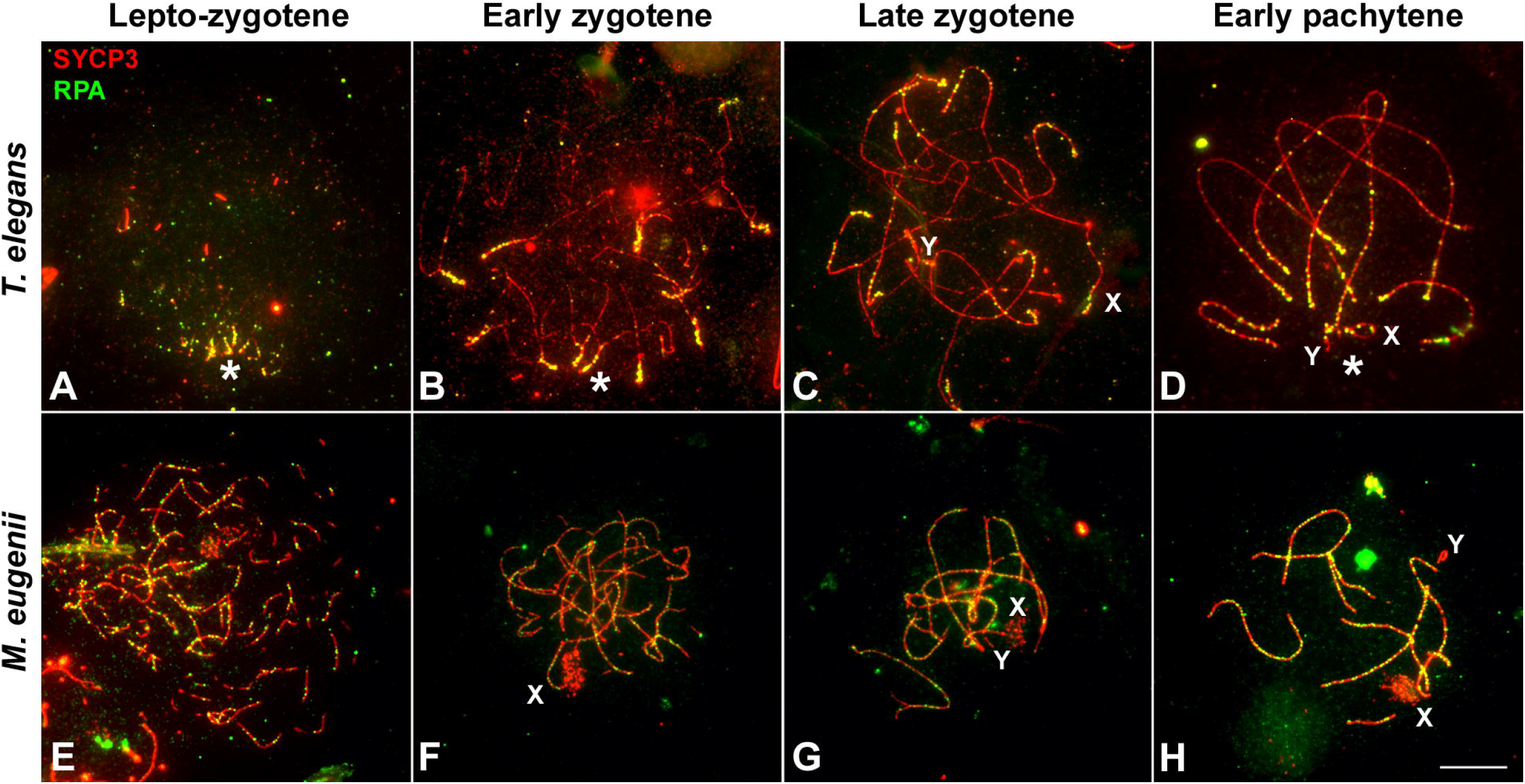
Localization of RPA. Spread spermatocytes labeled with antibodies against SYCP3 (red) and RPA (green). **A-D**. *T. elegans*. Most RPA foci concentrate in the distal regions of autosomes from leptotene to pachytene. From late zygotene to pachytene, RPA foci also accumulate on the sex chromosomes (X, Y). Asterisk indicates the polarization of chromosomal ends. sex chromosomes. **E-H**. *M. eugenii*. RPA foci associate to forming AEs at the leptotene to zygotene transition. From early zygotene onwards, foci are observed along the entire length of the bivalents. Bar: 10 μm.

### Chromosomal distribution of RPA

A remarkable feature observed regarding RPA dynamics was that the number of foci remained high even during early pachytene. Because autosomes have completed synapsis at this stage, every bivalent could be identified thanks to the differences in length, centromere position and location of the NOR (Figure 1B, 1D and Supplementary Figure 1). This permitted a quantitative study of the distribution of RPA along each chromosome in both *T. elegans* and *M. eugenii*. We analyzed at least 15 pachytene spermatocytes in two individuals from each species. Each bivalent was measured, divided into 10 segments and the position of each RPA focus was then scored along the bivalent and assigned to a segment. The same methodology was applied to the X chromosome for both species.

We detected that in *T. elegans* RPA foci accumulated towards the chromosomal ends in all bivalents, particularly in the four largest, in which the two distal segments concentrated near or above 50% of all RPA foci (Figure 7, Table 1). The distribution of RPA foci increased symmetrically in both chromosomal arms, with just a slight reduction around centromeres. This was especially relevant for bivalent 6, which bears the NOR on the short arm. This region accumulated fewer RPA foci (11.41%) compared to the opposite chromosomal end (20.16) (Table 1). The X chromosome, which remained as univalent, also showed a non-random distribution of RPA. The X centromere seemed to have an effect, with a reduced number of RPA in the flanking segments. The Y chromosome could not be analyzed due to its small size.

**FIGURE 7.**
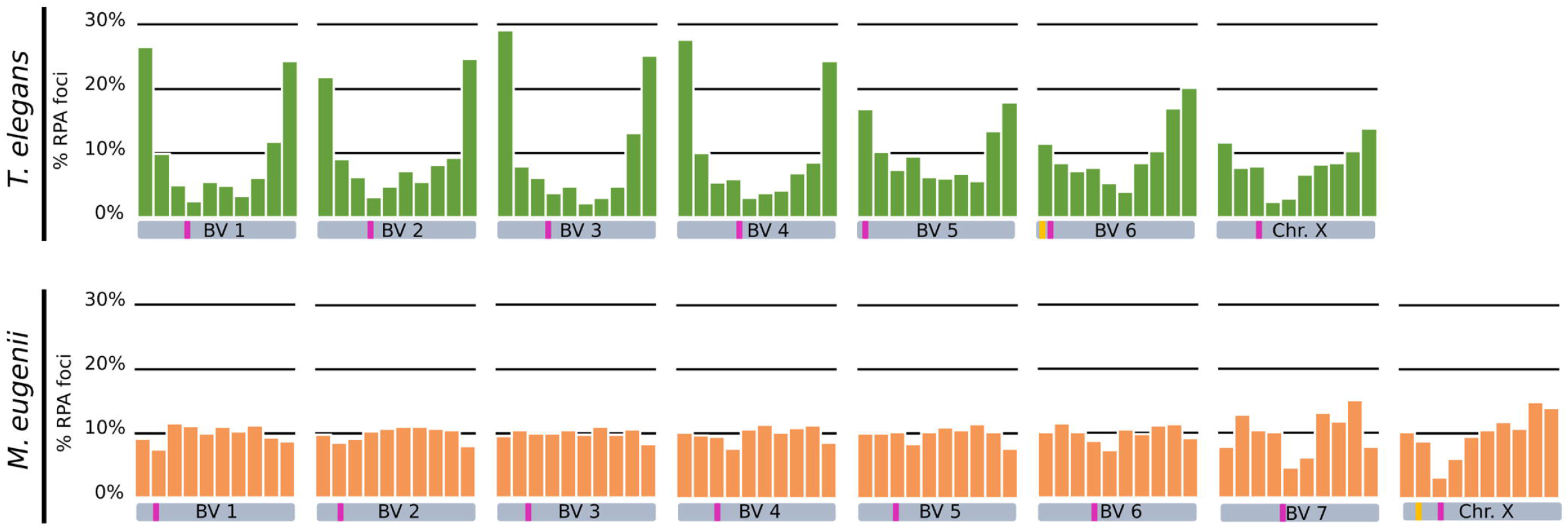
Chromosomal distribution of RPA foci distribution along bivalents and the X chromosome in *T. elegans* and *M. eugenii*. Chromosomes have been divided into 10 segments of equal size between telomeres (proximal to distal). Y axes in the graphs indicate the percentage of RPA foci in each segment. Each bivalent (BV) has been depicted below the corresponding graph. Pink bars indicate the centromere position, yellow bars indicate the NOR. We found a prominent polarization of RPA foci towards the chromosome ends in *T. elegans*. In contrast, in *M. eugenii* detection of RPA foci uncovered a relatively homogeneous distribution along the entire length of the chromosomes.

**Table 1.** Percentage of RPA foci per chromosomal segment in the different bivalents (BV) and the X chromosome (X) of T. elegans and M. eugenii. A χ^2^ of goodness of fit test (9 degrees of freedom) was performed to assess the deviation from a random distribution along chromosomes. Significance was considered when p≤ 0.05. n: number of bivalents analyzed.

| *Thylamys elegans* (% RPA) |  |  |  |  |  |  |  |  |
| --- | --- | --- | --- | --- | --- | --- | --- | --- |
| Segment | BV1<br>(n=31) | BV2<br>(n=35) | BV3<br>(n=33) | BV4<br>(n=35) | BV5<br>(n=33) | BV6<br>(n=35) | X<br>(n=28) |  |
| 1 | 26.51 | 21.85 | 29.10 | 27.64 | 16.87 | 11.41 | 14.57 |  |
| 2 | 9.90 | 9.07 | 7.88 | 10.00 | 10.24 | 8.49 | 9.60 |  |
| 3 | 5.03 | 6.30 | 6.13 | 5.45 | 7.43 | 7.16 | 9.93 |  |
| 4 | 2.52 | 3.15 | 3.72 | 6.00 | 9.44 | 7.69 | 2.98 |  |
| 5 | 5.54 | 4.81 | 4.81 | 3.09 | 6.22 | 5.31 | 3.64 |  |
| 6 | 4.87 | 7.22 | 2.19 | 3.82 | 6.02 | 3.98 | 8.28 |  |
| 7 | 3.36 | 5.56 | 3.06 | 4.18 | 6.83 | 8.49 | 10.26 |  |
| 8 | 6.21 | 8.15 | 4.81 | 6.91 | 5.62 | 10.34 | 10.60 |  |
| 9 | 11.74 | 9.26 | 13.13 | 8.55 | 13.45 | 16.98 | 12.91 |  |
| 10 | 24.33 | 24.63 | 25.16 | 24.36 | 17.87 | 20.16 | 17.22 |  |
| $\chi^2$ | 397.19 | 256.15 | 377.73 | 377.02 | 93.28 | 88.79 | 52.76 | |
| p value | <0.0001 | <0.0001 | <0.0001 | <0.0001 | <0.0001 | <0.0001 | 0.0004 |  |

| *Macropus eugenii* (% RPA) |  |  |  |  |  |  |  |  |
| --- | --- | --- | --- | --- | --- | --- | --- | --- |
| Segment | BV1<br>(n=43) | BV2<br>(n=44) | BV3<br>(n=44) | BV4<br>(n=44) | BV5<br>(n=43) | BV6<br>(n=43) | BV7<br>(n=42) | X<br>(n=31) |
| 1 | 9.18 | 9.80 | 9.59 | 10.11 | 10.02 | 10.17 | 7.87 | 10.29 |
| 2 | 7.54 | 8.56 | 10.49 | 9.67 | 10.02 | 11.50 | 12.79 | 8.82 |
| 3 | 11.54 | 9.13 | 9.99 | 9.56 | 10.27 | 10.17 | 10.49 | 3.19 |
| 4 | 11.08 | 10.37 | 9.99 | 7.69 | 8.42 | 8.83 | 10.16 | 6.13 |
| 5 | 9.97 | 10.75 | 10.49 | 10.66 | 10.27 | 7.33 | 4.59 | 9.56 |
| 6 | 11.02 | 11.04 | 9.79 | 11.43 | 11.01 | 10.50 | 6.23 | 10.54 |
| 7 | 10.30 | 11.04 | 10.99 | 10.11 | 10.52 | 9.83 | 13.11 | 11.76 |
| 8 | 11.28 | 10.75 | 9.79 | 10.88 | 11.51 | 11.17 | 11.80 | 10.78 |
| 9 | 9.38 | 10.56 | 10.59 | 11.32 | 10.27 | 11.33 | 15.08 | 14.95 |
| 10 | 8.72 | 7.99 | 8.29 | 8.57 | 7.67 | 9.17 | 7.87 | 13.97 |
| $\chi^2$ | 22.94 | 11.16 | 4.98 | 11.54 | 9.47 | 8.93 | 30.31 | 43.81 |
| p value | 0.006 | 0.2649 | 0.8357 | 0.2406 | 0.3946 | 0.4435 | 0.0004 | <0.0001 |

In contrast, the distribution of RPA foci in *M. eugenii* was quite homogeneous along bivalents. A χ^2^ test showed that on most chromosomes RPA location did not significantly depart from a random distribution (Table 1). The only exceptions were chromosomes 1, 7 and X, on which RPA foci were slightly reduced around the centromere. The NOR, which is located in the short arm of the X chromosome (Figure 1D) did not have and apparent effect on accumulation of RPA foci. In fact, RPA distribution was quite similar on the X chromosome in both species.

The quantitative analysis of RPA also allowed us to assess a potential correlation between the number of foci accumulated on every chromosome and their respective length. Because the X chromosome was a univalent and the Y chromosome was too small, we only considered autosomes. We found that in *T. elegans*, RPA foci were underrepresented in bivalents 1 to 3, and conversely overrepresented in bivalents 4 to 6 (Table 2). Accordingly, a Spearman correlation analysis of RPA foci number and SC length showed low correlation (r=0.41, p<0.0001) (Figure 8). In contrast, all *M. eugenii* bivalents presented an increased correlation between chromosome length and RPA proportion 11 (Spearman correlation analysis r=0.88, p<0.0001) (Table 2, Figure 8). This reinforces the hipothesis that RPA distribution in *M. eugenii* is not dependent on specific features of chromosomes. Their location was equiprobable on any chromosome and at any chromosomal region.

**Table 2.** Proportion of RPA foci and SC length of each bivalent at early pachytene, calculated over the number of foci and SC length of autosomes. Only cells in which all bivalents could be recorded have been included. Values are presented as mean ± standard deviation. n: number of cells analyzed.

| *Thylamys elegans* (n=34) |  |  |  |  |  |  |
| --- | --- | --- | --- | --- | --- | --- |
|  | BV1 | BV2 | BV3 | BV4 | BV5 | BV6 |
| % RPA foci | 19.57 ± 4.7 | 17.91 ± 6.7 | 15.98 ± 4.9 | 18.52 ± 7.0 | 15.20 ± 4.3 | 12.81 ± 4.9 |
| % SC length | 23.89 ± 1.7 | 21.89 ± 1.7 | 19.70 ± 1.9 | 15.12 ± 2.1 | 9.82 ± 1.3 | 8.80 ± 1.0 |

| *Macropus eugenii* (n=41) |  |  |  |  |  |  |  |
| --- | --- | --- | --- | --- | --- | --- | --- |
|  | BV1 | BV2 | BV3 | BV4 | BV5 | BV6 | BV7 |
| % RPA foci | 23.84 ± 3.9 | 16.81 ± 3.6 | 15.75 ± 3.1 | 14.50 ± 2.8 | 13.68 ± 3.1 | 10.49 ± 2.5 | 4.93 ± 2.0 |
| % SC length | 22.91 ± 1.9 | 16.96 ± 2.0 | 15.37 ± 1.5 | 14.47 ± 2.6 | 13.89 ± 1.4 | 10.93 ± 1.8 | 5.47 ± 0.9 |

**FIGURE 8.**
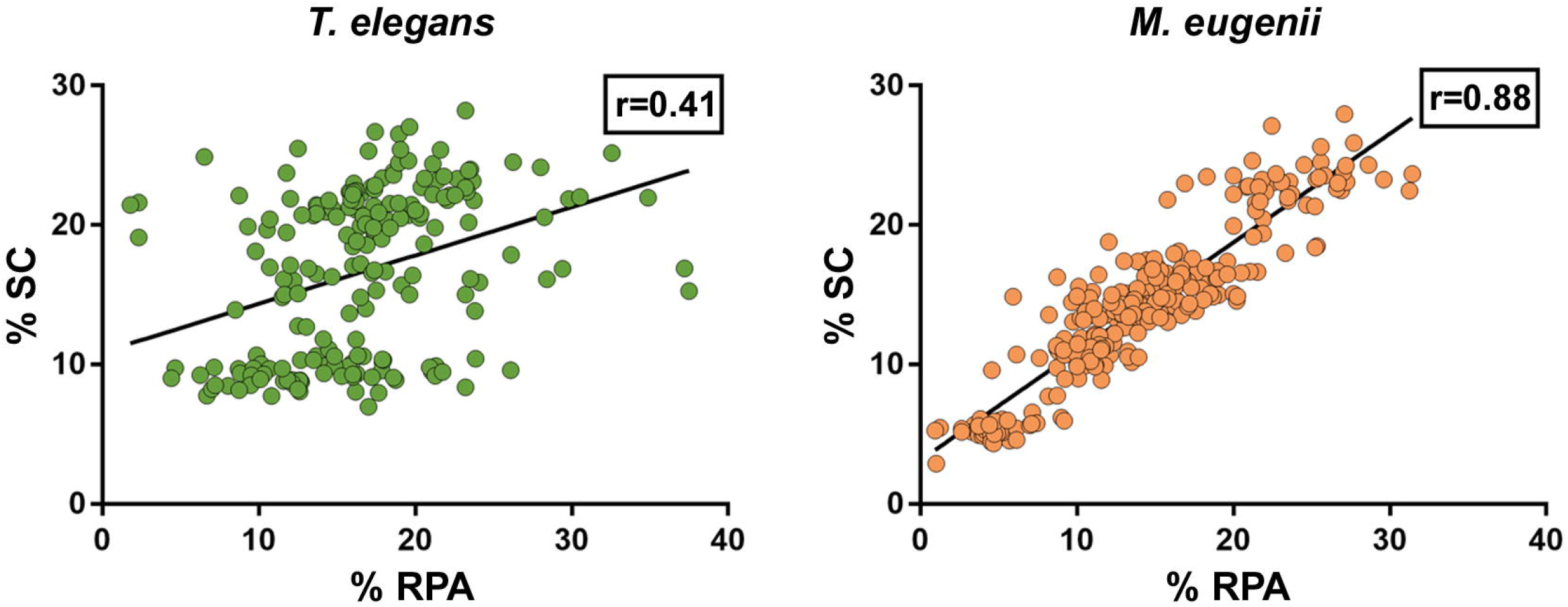
Compared proportion of RPA foci number and SC length chromosome in *T. elegans* and *M. eugenii*. Each spot represents a bivalent in a cell. Black lines represent the calculated regression line. r= Spearman correlation coefficient.

## DISCUSSION

Meiotic studies in non-eutherian mammalian species are scarce. Only a few reports were devoted to monotremes (Daish et al., 2015; Casey et al., 2017). In marsupials, most studies have focused on the unique behavior of sex chromosomes (Solari and Bianchi, 1975; Sharp, 1982; Roche et al., 1986; Seluja et al., 1987; Page et al., 2003; Page et al., 2005; Fernández-Donoso et al., 2010; Marín-Gual et al., 2022b). Our recent work on marsupials revealed divergent strategies for meiotic DNA repair, recombination and transcription (Marín-Gual et al., 2022b). Here we extend these observations and report previously uncharacterized features of marsupial meiosis: synapsis initiation and chromosomal distribution of DSBs. Remarkably, our observations suggest an evolutionary shift in the meiosis program between American and Australian marsupials. In the context of recently published reports on fish and reptile meiosis (Blokhina et al., 2019; Marín-Gual et al., 2022a), our results reveal the persistence of ancestral vertebrate meiotic features in marsupials. This highlights the relevance of comparative studies to fully understand the causes and consequences of meiosis evolution.

### The conspicuous *bouquet* conformation could be an ancient feature of vertebrate meiosis

The polarization of telomeres at the beginning of meiosis has been described in a wide range of species, from fungi to plants and animals (Zickler and Kleckner, 1998). However, the presence and the extent of this polarization changes from taxa to taxa, and even between sexes. Formation of the *bouquet* has been considered a crucial factor in chromosome pairing and synapsis initiation (Liebe et al., 2006; Reig-Viader et al., 2016). However, the study of different mutants has provided evidence that such a feature is not an absolute requirement, and some model systems like *Drosophila melanogaster* and *Caenorhabditis elegans* are known to lack a chromosomal *bouquet* during meiosis (Harper et al., 2004).

Our results show the presence of a *bouquet* in two of the three marsupial species analyzed. This was correlated with the initiation of synapsis, which was clearly terminal in *T. elegans* and *D. gliroides*. Synapsis in these two species extended from the telomeres to the interstitial regions in a zipper-like manner. Similar observations were reported in the South American marsupial *Rhyncholestes raphanurus*, belonging to the Paucituberculata Order (Page et al., 2005). Thus, this feature seems to be an old character among marsupials. Moreover, according to recent reports in zebrafish (Blokhina et al., 2019) and several species of reptiles (Marín-Gual et al., 2022a), these seem to be ancient features of the vertebrate meiotic program, which have been subsequently maintained in a wide range of groups. However, this chromosomal polarization has suffered regulatory modifications in different linages. In eutherian mammals, the house mouse displays a visible polarization at the beginning of meiosis (Berrios et al., 2010; Berrios et al., 2014), but this is brief and often incomplete (Scherthan et al., 1996; Lopez-Jimenez et al., 2022). Although synapsis can be initiated at the chromosomal ends, it can also start interstitially (Boateng et al., 2013). This is what we observed in the third marsupial species analyzed in this study, *M. eugenii*, which mimics the pattern described in mouse. Considering the phylogenetic relationships between the marsupials studied here (Figure 1) (Duchêne et al., 2017), and previous reports in other vertebrates (Blokhina et al., 2019; Marín-Gual et al., 2022a), the most parsimonious explanation is that *bouquet* polarization is an ancestral character in marsupials, and most probably in all vertebrates.

Consequently, the loss of telomere polarization at the beginning of meiosis seems to have occurred independently several times in the evolution of mammals (i.e., Australian marsupial species and mouse). What could cause this change in chromosome behavior? The determinants of *bouquet* polarization include the binding of telomeres to the nuclear envelope and their interaction with cytoskeleton components via transmembrane proteins of the nuclear envelope (Scherthan, 2007). It is unlikely that these dynamics have been lost in eutherian mammals or in Australian marsupials, but they could be regulated differently. Interestingly, a recent report revealed that the formation of a primary cilium in spermatocytes is a crucial factor in the formation of the *bouquet* in zebrafish (Mytlis et al., 2022). This structure is formed in spermatocytes at leptotene-zygotene, and its removal disrupts *bouquet* formation, as well as synapsis and recombination (Mytlis et al., 2022; Xie et al., 2022). Intriguingly, the formation of the primary cilium in mouse seems to be differentially regulated, because this structure is formed in a reduced fraction of spermatocytes at the leptotene-zygote transition (Lopez-Jimenez et al., 2022). This seems to correlate with the absence of a conspicuous *bouquet* in the mouse, although a causative relationship has not yet been demonstrated. Further exploration of mammalian species could provide new insight into the role that cilia play in *bouquet* formation.

### Induction of DSBs and synapsis initiation and progression

One of the most characteristic hallmarks of meiotic DNA damage is the localization of the phosphorylated form of H2AX (γH2AX) (Mahadevaiah et al., 2001; Turner et al., 2004), which appears as scattered foci at early leptotene and then extends to occupy the whole nucleus during late leptotene (Enguita-Marruedo et al., 2019). In eutherian mammals (i.e., mouse) the presence of γH2AX is then reduced as prophase I progresses and DNA is repaired but remains during late stages of prophase I in regions that do not achieve synapsis. This has been found to occur on both autosomes, as a feature related to the meiotic silencing of unsynapsed chromatin (MSUC) (Baarends et al., 2005; Turner et al., 2005; Manterola et al., 2009), and on the sex chromosomes where it contributes to meiotic sex chromosome inactivation (MSCI) (Turner et al., 2004; Page et al., 2012). Early reports on the localization of γH2AX in marsupials indicated that MSCI also operates in this group (Franco et al., 2007; Hornecker et al., 2007; Namekawa et al., 2007). However, other aspects of the localization of γH2AX in relation to DNA damage in marsupials have remained unexplored until recently.

We have previously shown in marsupials that there are two waves of γH2AX accumulation during prophase I, along with lower levels of γH2AX on autosomes when compared to eutherians (Marín-Gual et al., 2022b). Here we extend these initial observations and report previously uncharacterized differences between marsupial species. In *T. elegans* and *D. gliroides* γH2AX signal is scarce and restricted to the regions where homologous chromosomes initiate their synapsis, whereas in *M. eugenii* γH2AX is distributed across the whole nucleus. We suggest that there is a relationship between this finding and the observed patterns of synapsis progression. Thus, in the two American species a low induction of DSBs would occur in the chromosomal regions polarized to the *bouquet* area, triggering the initiation of synapsis. In the absence of further (or abundant) DSBs along interstitial regions of chromosomes, synapsis would progress from chromosomal ends towards the center of chromosomes, probably owing to the self-assembly capabilities of the SC components. Therefore, the few DSBs scattered along interstitial regions do not seem to promote SC assembly. Interestingly, these interstitial DSBs do not trigger a conspicuous H2AX phosphorylation either. Only later, during late zygotene, is γH2AX observed at interstitial regions of the unsynapsed autosomes, as well as on the sex chromosomes. This could be interpreted as an indication of late DNA damage events produced exclusively in those regions. Alternatively, it might be linked to the silencing of unsynapsed regions, i.e., the MSUC/MSCI processes. In contrast, the widespread generation of DSBs in *M. eugenii* is correlated to synapsis initiation at different regions along the chromosomes, not only chromosomal ends. Thus, the synapsis pattern of homologous chromosomes seems to be conditioned by the way DNA damage is produced during prophase I. The pattern observed in *T. elegans* and *D. gliroides* seems to be ancestral, and even shared by other non-mammalian vertebrates (Blokhina et al., 2019; Marín-Gual et al., 2022a).

Finally, we highlight the possibility that a part of the DNA damage occurring in *T. elegans* and *D. gliroides* was not accompanied by H2AX phosphorylation. Although the restricted localization of γH2AX at the *bouquet* area correlates with accumulation of RPA and RAD51 in these two species, some RPA and RAD51 foci appeared outside the areas of γH2AX accumulation. Previous reports in monotreme mammals (Daish et al., 2015) and some insects (Viera et al., 2017) have indicated that γH2AX is not necessarily a marker of DNA damage during prophase I. Our own observations indicate that γH2AX is not detected during prophase I in some reptiles (Marín-Gual et al., 2022a) (Page, unpublished). Therefore, is seems that some aspects of DNA damage signaling during meiosis in mammals and other vertebrates are yet to be properly characterized.

### Differential chromosomal distribution of meiotic DSBs in marsupials

Perhaps the most striking finding of this study is the extreme difference in the distribution of DNA damage along chromosomes in the species analyzed. Several studies have focused on the overall frequency of recombination across mammals or even eukaryotes (Dumont and Payseur, 2008; Segura et al., 2013; Stapley et al., 2017). In eutherian mammals, previous reports have found that early diverging linages had lower recombination rates (Segura et al., 2013). Furthermore, marsupials show an even lower rate of recombination when compared to eutherians (Zenger et al., 2002; Samollow et al., 2004; Samollow et al., 2007; Dumont and Payseur, 2008; Wang et al., 2011), which has been attributed to the induction of fewer DSBs during early stages of meiotic prophase I (Marín-Gual et al., 2022b).

While many studies have stressed the evolutionary relevance of the recombination rate on chromosomal evolution and populations dynamics (Farré et al., 2012; Capilla et al., 2014; Ullastres et al., 2014; Dapper and Payseur, 2017; Ritz et al., 2017), the genomic implications of the uneven distribution of recombination along chromosomes have received less attention. Initial reports in mouse and human showed that COs tend to locate towards the telomeres (Barlow and Hultén, 1998; Froenicke et al., 2002). Likewise, studies on the localization of DSBs by means of DNA repair markers like RPA reported an accumulation of breaks towards the chromosomal ends (Oliver-Bonet et al., 2007; Pratto et al., 2014). Here we reveal striking inter-specific differences in the pattern of RPA distribution (CO distribution could not be analyzed due to the lack of reactivity of many different antibodies against MLH1 and other CO markers) in marsupials. In *T. elegans* RPA foci accumulated towards the chromosomal ends. This pattern resembles the one characterized in eutherian mammals, although it is more prominent in marsupials. In contrast, *M. eugenii* shows a remarkably even distribution of DSBs along the chromosomes. Moreover, the number of DSBs per chromosome has a high correlation with SC (Table 2, Supplementary Figure 2), while in *T. elegans* large chromosomes appear to accumulate less RPA foci than expected.

We can only speculate on the mechanisms and consequences of the differential chromosomal distribution of DSBs detected in marsupials. In mammals, and many other organisms, DSBs are produced preferentially at recurrent sites referred to as hotspots (Paigen and Petkov, 2010; Baudat et al., 2013). Two main types of hotspots are usually recognized. The first are placed in promoter regions of genes, which supposedly present an open chromatin configuration that makes them accessible to the DSBs producing complexes. The second type is determined by the action of the histone methyl transferase PRDM9, which tri-methylates histone H3 at lysine 4 (H3K4me), thus transforming these sites into preferential spots for breakage by the protein SPO11 (Baudat et al., 2010; Parvanov et al., 2010; Brick et al., 2012). Whereas the first kind of hotspots are conserved within and between species, the ones depending on PRDM9 are more variable, owing to the fast-evolving features of this enzyme (Grey et al., 2018). Most mammals, including marsupials have a copy of the *Prdm9* gene, but it has been partially or completely lost in the platypus and canids (Cavassim et al., 2022). Therefore, it seems unlikely that the different patterns of DSB distribution we observe could be attributable to the absence of PRDM9. Instead, it is possible that a differential distribution, usage, or regulation of the different types of hotspots could be responsible for such differences. One possibility is that *T. elegans* and *D. gliroides* rely more in the use of promoter-related hotspots, with genes concentrated near chromosomal ends.

*M. eugenii* could be using more PRDM9-dependent hotspots, which would be expected to result in a more uniform distribution of DSBs along chromosomes. This even distribution in *M. eugenii* could be also related to the extensive genomic reorganizations experienced in the family Macropodidae (Deakin, 2018; Deakin and O’Neill, 2020; Álvarez-González et al., 2022). In fact, recent reports have shown that lineage-specific evolutionary genomic reshuffling can influence patterns of higher-order chromatin organization (Farré et al., 2015; Álvarez-González et al., 2022), and that chromosomal reorganizations can have an impact on the three-dimensional genome folding and recombination in the germ line (Vara et al., 2021; Álvarez-González et al., 2022). Thus, genome reshuffling in macropodids could have led to a more even distribution of recombination hotspots genome-wide. Interestingly, we found that the X chromosome behaves similarly in the species compared here. It is possible that the X chromosome escaped this hotspot reorganization. Further analyses in the study of marsupial genomes and the use of ChIP-Seq approaches to map recombination hotspots could yield insightful information about this possibility.

### Genomic and evolutionary consequences of divergent recombinogenic patterns

The dissimilar pattern of DSBs chromosomal distribution may have consequences at the genomic level in marsupials. Although not all the DSBs produced during prophase I result into COs, it seems reasonable to assume that COs could be evenly distributed along chromosomes in *M. eugenii*. This would facilitate the recurrent recombination of allele combinations, thus breaking haplotypes. In the case of *T. elegans* and *D. gliroides*, however, the accumulation of DSBs towards chromosomal ends would reduce the possibilities of recombination at the interstitial regions of chromosomes. Supporting this view, a previous study reported that chiasmata are conspicuously terminal in *T. elegans* (Page et al., 2006). Since both *T. elegans* and *D. gliroides* have a very low chromosome number, this would mean the formation of few and large regions of linkage disequilibrium. Such a strategy could be beneficial in a very stable environment, as long as allelic combinations at different loci had achieved an optimum (Stapley et al., 2017; Wang et al., 2019). In contrast, with recombination spread all over chromosomes, the resulting generation of new genome-wide allele combinations could have provided some marsupial groups with a higher capacity to adapt to new environments. It is tempting to speculate that this factor could have had an influence in the diversification of marsupials in Australasia after they diverged from Microbiotheria.

Moreover, the fact that most DSBs are repaired as gene conversion events (Cole et al., 2012; Baudat et al., 2013) does not preclude these breaks from being innocuous for the evolution of some genome features, like GC content. Both reciprocal recombination and gene conversion induce a shift to the accumulation of GC in the repaired strand, a phenomenon known as GC-biased gene conversion (gBGC) (Duret and Galtier, 2009). This mechanism has been detected from yeast to mammals and has been proposed to impact the evolution of genomes (Mugal et al., 2015). For instance, it was suggested that the enrichment of GC-rich isochores in mammalian genomes could be in part a consequence of gBGC (Duret and Galtier, 2009). The accumulation of GC content due to gBGC requires the recurrent use of recombination hotspots. Given the differential use of these hotspots across species, different rates of GC accumulation are expected. This could partially explain why in humans, with a rapid turnover of PRDM9-dependen hotspot, GC accumulation is spread in the whole genome, whereas in birds, with more conserved recombination hotspots, the increase of GC content is much more localized to specific genomic regions (Mugal et al., 2015). Thus, given the distribution of DSBs in the marsupial species studied, we foresee an accumulation of GC content due to gBGC at the distal regions of chromosomes in *T. elegans* and *D. gliroides*, compared to *M. eugenii*.

### Concluding remarks

Our results suggest that marsupials experienced a major shift in some of the key processes of meiosis, such as SC assembly, synapsis progression and DSBs distribution (Figure 9). This shift seems to have occurred after the split of Microbiotheria and the Australian marsupials, about 60 million years ago (Feng et al., 2022). However, to what extent the features observed in *M. eugenii* are shared by other Australian marsupials needs further validation. Likewise, the features observed in *T. elegans*, clearly basal to the rest of the marsupial groups, could have been shared with the ancestor of the eutherian mammals before they split apart about 165 million years ago. In view of recent reports, these features could be even dated back to the appearance of early vertebrates (Blokhina et al., 2019; Marín-Gual et al., 2022a). Expansion of meiosis studies to uncharacterized mammals, including eutherians, marsupials and monotremes, as well as to other vertebrates (i.e., reptiles, amphibians or fishes), will shed light on the evolution of meiosis across taxa. Moreover, these studies will undoubtedly have a deep impact in our understanding of genome evolution.

**FIGURE 9.**
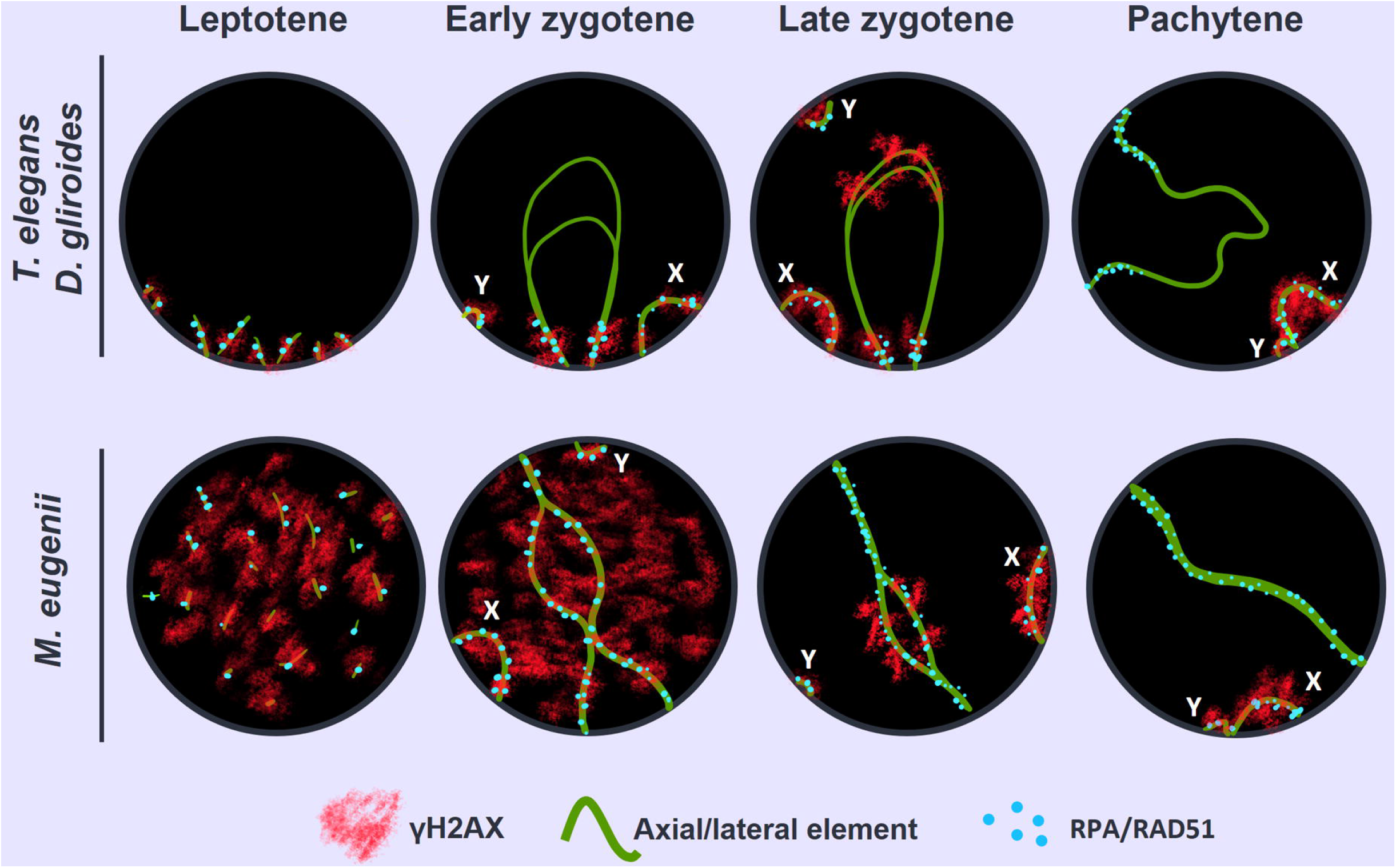
Schematic representation of the two different patterns of SC assembly, synapsis progression and DNA repair observed in the marsupial species. Green lines represent AE/LEs, red clouds represent γH2AX and blue spots represent DNA repair proteins (RAD51/RPA). In each nucleus a bivalent and both sex chromosomes (X, Y) are represented. In the top row, the putative ancestral pattern involves: conspicuous *bouquet* polarization, AE assembly and synapsis progression from chromosomal ends and preferential localization of DNA damage and repair towards chromosomal distal regions. In the bottom row, the emergent pattern observed in Australian marsupials: loosened *bouquet*, AEs assembly and synapsis progression at any chromosomal position, and an even distribution of DNA damage and repair events along chromosomes.

## ACKNOWLEDGEMENTS

This work was supported by grants CGL2014-53106-P to J.P. (Ministerio de Ecomonía y Competitividad, Spain), BIOUAM02-2020 to J.P. and R.G. (Departamento de Biología, Univerdiad Autónoma de Madrid), PID2020-112557GB-I00 to A. R.-H. (Ministerio de Ciencia e Innovación, Spain) and from the Australian Research Council to M.B.R., G.S. and P.D.W. (DP21103512 and DP220101429). P.D.W is also supported by the NHMRC (APP1182667 and APP2021172). L.M.-G. was supported by a FPU predoctoral fellowship from the Ministry of Science, Innovation and University (FPU18/03867). We are indebted to Dr. Paula Cohen, Dr. Alberto Pendas, Dr. Monica Pradillo, Dr. Jose A. Suja and Dr. Attila Toth for sharing antibodies against crossover markers. We thank Corinne van Den Hoek and members of the wallaby research group (Walgroup) for assistance with the wallabies. We dedicate this work to the loving memory of Prof. Francisco Bozinovic, for his enthusiasm and motivation in the study of Chilean fauna.

---

SUPPLEMENTARY FIGURE 1. Identification of bivalents. Spread spermatocytes at pachytene labeled with antibodies against SYCP3 (red), RPA (green), centromere proteins (blue) and fibrillarin (pink). **A**. *T. elegans*. **B**. *M. eugenii*. The length and centromere position allowed for the identification of each bivalent. Additionally, bivalent 6 in *T. elegans* was recognized by the position of the nucleolus (Nu), revealed by fibrillarin labeling. Sex chromosomes (X, Y).

