## Supplementary figures and images for "Divergent patterns of meiotic double strand breaks and synapsis initiation dynamics suggest an evolutionary shift in the meiosis program between American and Australian marsupials"

### Supplementary Figure 1

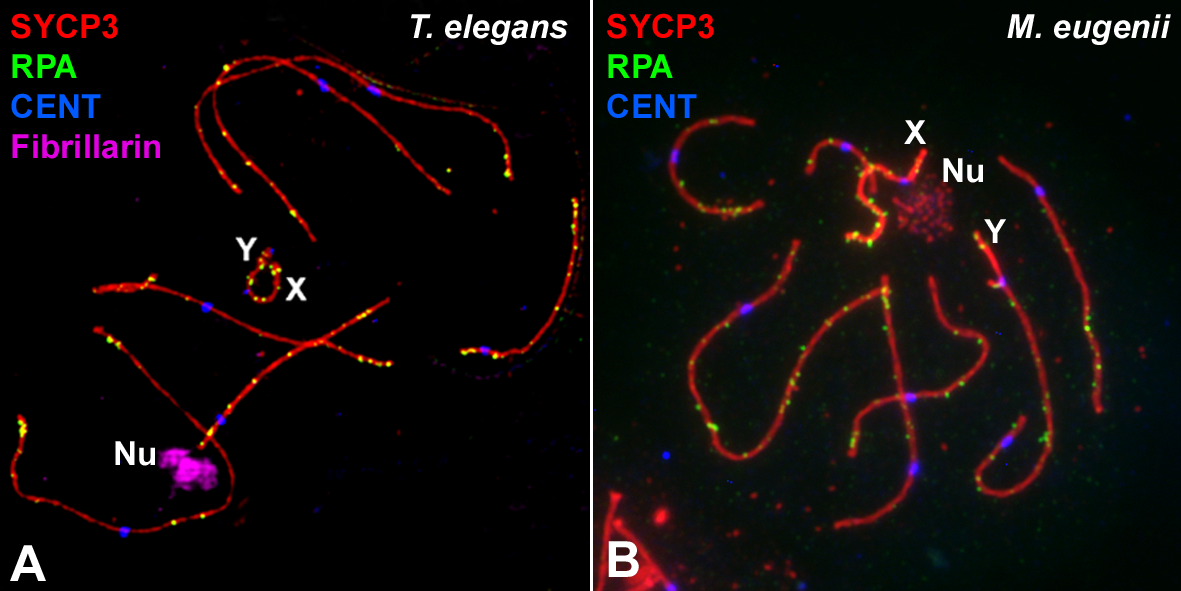
